# Stone Age “chewing gum” yields 5,700 year-old human genome and oral microbiome

**DOI:** 10.1101/493882

**Authors:** T. Z. T. Jensen, J. Niemann, K. Højholt Iversen, A.K. Fotakis, S. Gopalakrishnan, M.-H.S. Sinding, M. R. Ellegaard, M. E. Allentoft, L. T. Lanigan, A. J. Taurozzi, S. Holtsmark Nielsen, M.W. Dee, M. N. Mortensen, M. C. Christensen, S. A. Sørensen, M. J. Collins, M.T.P Gilbert, M. Sikora, S. Rasmussen, H. Schroeder

## Abstract

We present a complete ancient human genome and oral microbiome sequenced from a piece of resinous “chewing gum” recovered from a Stone Age site on the island of Lolland, Denmark, and directly dated to 5,858-5,661 cal. BP (GrM13305;5,007±11). We sequenced the genome to an average depth-of coverage of 2.3× and find that the individual who chewed the resin was female and genetically more closely related to western hunter gatherers from mainland Europe, than hunter gatherers from central Scandinavia. We use imputed genotypes to predict physical characteristics and find that she had dark skin and hair, and blue eyes. Lastly, we also recovered microbial DNA that is characteristic of an oral microbiome and faunal reads that likely associate with diet. The results highlight the potential for this type of sample material as a new source of ancient human and microbial DNA.

---

Our ability to recover ancient genomes from archaeological bones and teeth has revolutionised our understanding of human prehistory (1). The recovery of ancient microbial DNA, meanwhile, has provided fascinating insights into human health in the past and the evolution of the human microbiome (2). However, such analyses are highly dependent on the preservation of suitable samples, which get increasingly scarce as we move back in time. Further, there are periodic gaps in the burial record that can be either explained by poor preservation or burial practices that did not leave any traces behind, e.g. cremations (3, 4). Considering the scarcity of human remains, we hypothesised that there might be other sources of ancient human DNA in the archaeological record, in particular ancient “chewing gum”. These inconspicuous lumps of birch tar have been recovered from several prehistoric sites in Scandinavia and elsewhere in Europe and while their use is still debated, they often show tooth imprints, suggesting that they were chewed (5).

Birch bark resin has been used as adhesive as far back as the Middle Pleistocene (6, 7). However, direct evidence of it being used as a masticant, first appears in the Early Holocene (5). The oldest pieces of chewed birch bark resin found in archaeological contexts, have been dated to the Preboreal of Northern Europe (c.11,700-9,000 BP), such as Huseby Klev in Sweden (8), and Barmose in Denmark (9). Several of these objects were subsequently identified as birch bark tar by gas chromatography-mass spectrometry (GC-MS) analysis, through detection of both triterpenoids betulin and lupeol, which when found together are unique to birch (*Betula*) (5). Both of these triterpenoids have medicinal properties, such as being anti-microbial, anti-viral, and anti-inflammatory, suggesting that the use of birch resin as masticant in prehistory may also have been medicinal (5).

We used next-generation sequencing to recover a complete ancient human genome from one such artefact (Fig. 1a) that was found at the Stone Age site of Syltholm in Denmark, and directly dated to 5,858-5,661 cal. BP (GrM-13305; 5,007±11) (Fig. 1b). The site is one of the largest Stone Age settlements excavated to date (10) and spans the transition from the Mesolithic to the Neolithic period in Denmark (Fig. S3). In addition to the human genome, we also recovered microbial reads from the gum that enabled us to reconstruct a 5,700 year-old oral microbiome. The human genome sheds light on the dynamics of European populations at the time of the Neolithic transition, while the microbial data highlight the effects of major dietary shifts associated with the introduction of agriculture on our ancestral oral microbiome. Additionally, we found reads that are likely associated with diet. Overall, the results of our study highlight the potential of birch tar mastics as a new and well preserved source of ancient DNA.

**Fig. 1.**
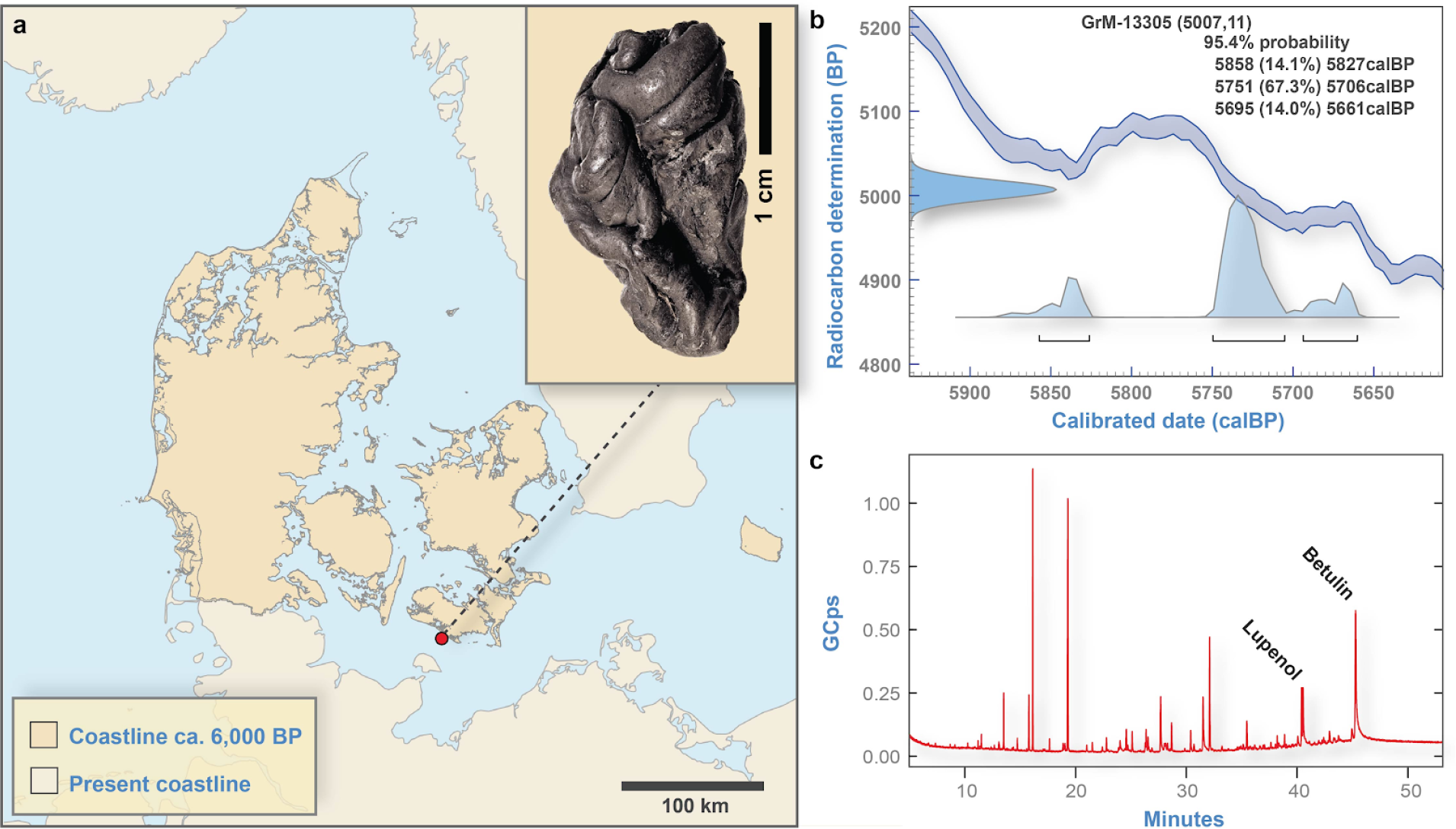
**a)** Photograph of the Syltholm “gum” and its find location at the site of Syltholm on the island of Lolland, Denmark (Danish coastline around 6,000 BP after ref. (87)); **b)** Calibrated date for the resin (5,858-5,661 cal. BP; 5,007 ±7); **C)** GC-MS chromatogram of the Syltholm resin showing the presence of triterpenes betulin and lupeol, which are characteristic for birch (*Betula pendula*) tar.

We generated over 380 million reads for this sample, over half of which mapped to the human reference genome (Table S2). We used the human reads to reconstruct the complete genome of the Syltholm individual, assembled to an effective depth-of-coverage of 2.3×. All the libraries displayed features characteristic for ancient DNA, including i) short average fragment lengths, ii) an increased occurrence of purines before strand breaks, and iii) an increased frequency of apparent cytosine (C) to thymine (T) substitutions at 5′-ends (Fig. S6). The amount of modern human contamination was estimated to be around 1-3% based on mtDNA (11). The sex of the individual could be determined to be female, based on the fraction of high quality reads (MAPQ ≥30) mapping to the sex chromosomes (12). In addition, we recovered a complete mitochondrial genome, sequenced to 91-fold coverage. The mtDNA haplogroup was determined to be K1e.

## Genome-wide affinities

To investigate the population genetic affinities of the Syltholm individual, we merged the ancient genome with over 1,200 modern and ancient genomes (Dataset S1), and performed a principal component analysis (PCA), by projecting the ancient sample onto the reference panel. We find that the Syltholm genome clusters with Palaeolithic and Mesolithic hunter-gatherers from western Europe, including Loschbour (13) and La Braña (14) (Fig. 2a). To explore the ancestral composition of the Syltholm genome further, we performed a supervised ADMIXTURE (15) run, specifying western hunter-gatherers (WHG), eastern hunter-gatherers (EHG), and Neolithic farmers (Barcın) as possible source populations. The result shows that the Syltholm genome is entirely composed of western hunter-gatherer ancestry, without any significant trace of EHG or farmer ancestry (Fig. 2b). To formally test these relationships, we computed a series of *f*- and *D*-statistics and found no evidence for significant levels of EHG or farmer gene flow in the Syltholm genome (Fig. 2c, Table S4-5). When using the modeling approach implemented in *qpAdm* (16), we find that a model assuming 100% WHG ancestry cannot be rejected in favour of two-way admixture. Specifically, modelling the Syltholm genome as a mixture of EHG and WHG ancestries does not significantly improve the fit, and neither does the addition of Neolithic farmer ancestry (Table S6).

**Fig. 2.**
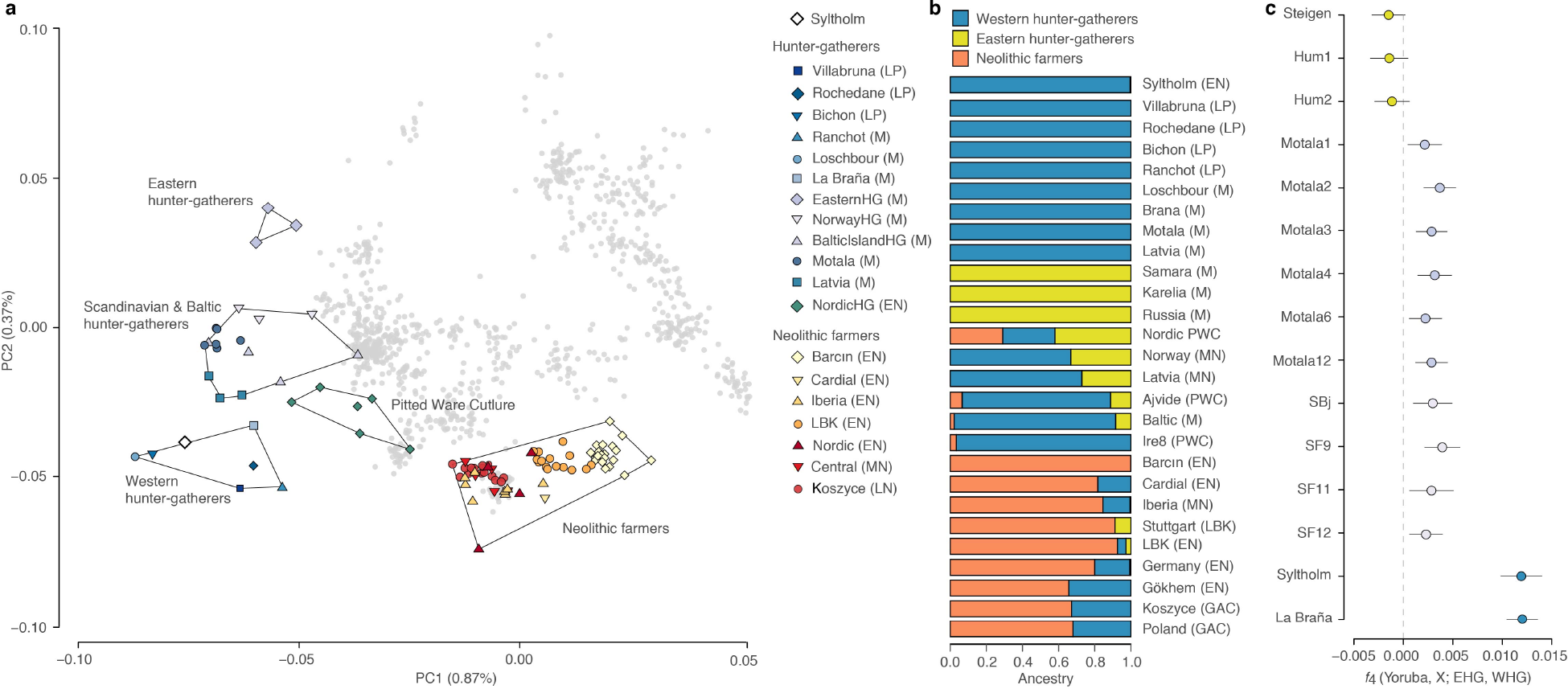
Genetic affinities of the Syltholm genome. **a)** PCA of modern Eurasian individuals (in grey) and a selection of ancient genomes, including the Syltholm genome. The ancient individuals were projected on the modern variation, **b)** Admixture proportions based on supervised ADMIXTURE analysis (15), specifying WHG, EHG, and Neolithic farmers (Barcın) as potential ancestral source populations, **c**) Allele-sharing between the Syltholm individual, Scandinavian hunter-gatherers, Baltic hunter-gatherers and EHG versus WHG measured by the statistic *f*_4_(Yoruba, *X*; EHG, WHG) where *X* stands for one of 15 ancient hunter-gatherer genomes.

## Pigmentation, lactase persistence and immune response

We used imputed genotypes for 41 SNPs included in the HIrisPlex-S system (17) to predict hair, eye and skin colour and found that the Syltholm female likely had blue eyes, brown hair and dark skin (Dataset S2). We also examined the allelic state of two SNPs linked with the primary haplotype associated with lactase persistence in modern humans and found that the she carried the ancestral allele for both, indicating that she was lactose intolerant (Dataset S3). In addition, we imputed alleles at 62 loci on 42 genes involved in the immune response in humans and found that she carried at least one derived allele in 21 (50%) of the genes involved (Dataset S3).

## Microbiome characterisation

Metagenomic profiling of the non-human reads in our sample revealed a close affinity between the microbial composition of the gum and the oral microbiome profiles from the Human Microbiome Project (HMP) (Fig. 3a). The actual profile (Fig. 3b) closely matches the profiles of oral microbiomes in HMP with two exceptions: a higher count of *Neisseriales* and a lower occurrence of *Veillonellales*. To further characterise and validate the microbial taxa present in our sample, we used MGmapper (18). As expected, the majority of reads could be assigned to taxa associated with the oral microbiome, with eight taxa having an effective depth-of-coverage >3× (Table 1). Additionally, we recovered 539 reads that mapped to Human herpes virus 4 (Epstein-Barr virus). We validated these taxa by aligning the non-human reads to the respective reference genomes and inspecting post-mortem DNA damage patterns (Fig. S7) and checking for evenness of coverage (Fig. S8).

**Fig. 3.**
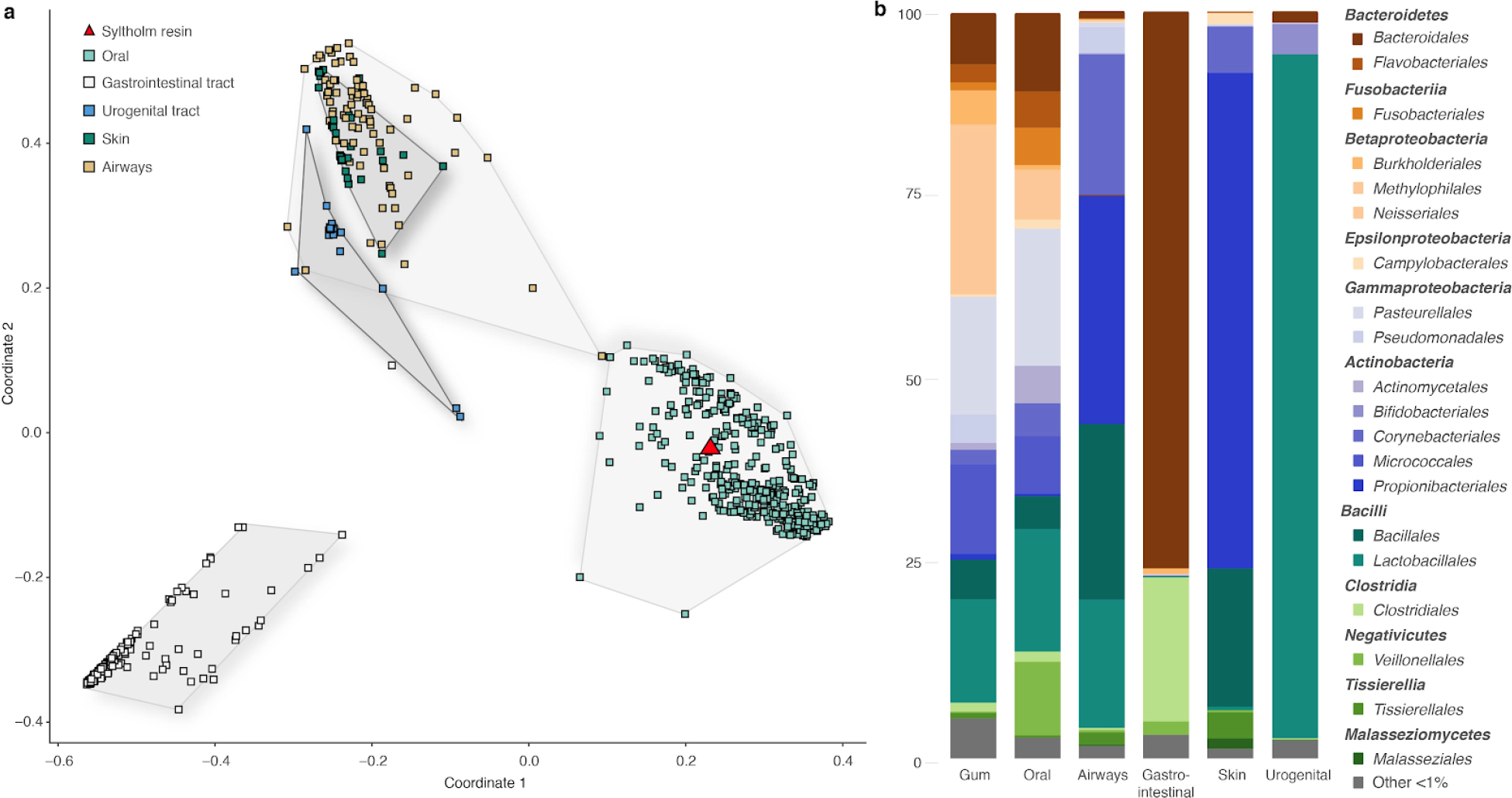
Metagenomic profile of the Syltholm “gum”. **a)** PCoA with Bray-Curtis at genera level with 689 microbiomes from HMP (79) and the Syltholm sample; **b)** Order-level microbial composition of the Syltholm sample compared to modern microbial profiles from HMP (79) visualised using MEGAN6 (80).

**Table 1.** Non-human taxa identified from metagenomic data recovered from the Syltholm sample.

| Species | Reads MQ35 | Av. read length (bp) | DoC | > 1X (%) | Delta S |
| --- | --- | --- | --- | --- | --- |
| *Actinomyces naeslundii* | 44,551 | 43 | 0.63 | 38.9% | 0.79 |
| *Actinomyces odontolyticus* | 19,829 | 41 | 0.35 | 23.4% | 0.62 |
| *Actinomyces viscosus* | 50,621 | 43 | 0.71 | 42.3% | 0.76 |
| *Fusobacterium periodonticum* | 28,822 | 63 | 0.82 | 43.5% | 0.45 |
| *Haemophilus haemolyticus* | 234,479 | 58 | 7.25 | 86.5% | 0.50 |
| *Neisseria cinerea* | 371,703 | 53 | 10.62 | 92.7% | 0.55 |
| *Neisseria meningitidis* | 167,449 | 49 | 3.68 | 51.6% | 0.45 |
| *Porphyromonas gingivalis* | 18,344 | 54 | 0.42 | 32.8% | 0.60 |
| *Prevotella intermedia* | 56,081 | 55 | 1.17 | 58.7% | 0.55 |
| *Rothia mucilaginosa* | 271,064 | 49 | 5.88 | 77.8% | 0.61 |
| *Streptococcus infantis* | 113,743 | 56 | 3.06 | 54.6% | 0.40 |
| *Streptococcus mitis* | 306,887 | 57 | 8.25 | 76.7% | 0.40 |
| *Streptococcus oralis* | 141,250 | 54 | 3.99 | 66.7% | 0.37 |
| *Streptococcus pneumoniae* | 256,392 | 57 | 7.20 | 72.5% | 0.39 |
| *Treponema denticola* | 13,968 | 57 | 0.28 | 22.1% | 0.44 |
| *Veillonella parvula* | 1,870 | 52 | 0.05 | 4.0% | 0.56 |
| *Human herpesvirus 4* | 539 | 49 | 0.15 | 13.6% | 0.60 |
| *Anas platyrhynchos* | 476,458 | 53 | 0.02 | 1.9% | 0.52 |
| *Anguilla anguilla* | 56,439 | 42 | 0.002 | 0.2% | 0.42 |
| *Betula pendula* | 25,343 | 54 | 0.003 | 0.3% | 0.49 |

## Diet

We also identified reads that could associated with taxa related to diet, including mallard (*Anas platyrhynchos*) and eel (*Anguilla anguilla*), as well as reads associated with the source of the resin: birch (*Betula pendula*) (Table 1). We validated the presence of these taxa, as described above, by checking for the presence of damage patterns (Fig. S7) and evenness of coverage (Fig. S8). We also specifically looked for reads mapping to other taxa, such as barley (*Hordeum vulgare*), emmer (*Triticum dicoccum*) and einkorn (*Triticum monococcum*), as well as common hazel (*Corylus avellana*), but either did not retain any reads after filtering for mapping quality (MAPQ ≥35) or were unable to confirm that the reads are ancient.

The results highlight the potential of birch resin mastics as a new source of ancient human and microbial DNA. The endogenous human DNA content was surprisingly high and the overall preservation of the DNA in the sample was comparable to well-preserved teeth or petrous bones (Table S3) (19–21). We believe that the remarkable preservation can be explained by the antiseptic, hydrophobic environment of the resin, which both inhibits microbial and chemical degradation of DNA. This opens up fascinating new possibilities to extend the spectrum of human ancient DNA studies, particularly for time periods or cultural contexts where human remains are scarce or non-existent. Furthermore, our ability to recover authentic microbial DNA provides information on the state and evolution of the human oral microbiome and the spread of specific oral pathogens, while reads that likely relate to foodstuffs provide fascinating insights into the diet and changing subsistence strategies of prehistoric people.

## Population history

From a population history point of view, the 2.3× Syltholm genome (5,858-5,661 cal. BP) provides an important data point at a crucial juncture in European prehistory, the time of Mesolithic/Neolithic transition. In Denmark, this transition occurred around 6,000 years BP and is marked by rapid and dramatic changes in diet and subsistence strategies from a largely marine diet in the Mesolithic to a terrestrial-based diet in the Neolithic (22–24). Culturally, the change is marked by the transition from the Late Mesolithic Ertebølle culture (7,300-5,800 cal. BP) with its flaked stone tools and typical T-shaped antler axes (25, 26), to the early Neolithic Funnel Beaker culture (5,900-5,300 cal. BP) with its characteristic pottery, polished flint artefacts, and domesticated animals (27–29). While human remains from Denmark exist for this time period, none have been sequenced to date. In fact, there are still only a handful of published ancient genomes from Scandinavia as a whole (30–33).

Previous studies have shown that the late hunter-gatherer populations of Europe can be split into western hunter-gatherers (WHG), typified by individuals from present-day Luxembourg (13) and Spain (14), and eastern hunter-gatherers (EHG), generally represented by individuals from northern Russia and Estonia (16). Genetically, hunter-gatherers from central Scandinavia are known to be intermediate along this cline, mirroring their geographic location between the two extremes of this west–east axis (13, 31). According to Günther et al. (32), EHG ancestry reached central Scandinavia from the north, while the WHG ancestry entered from the south, following the retreat of the ice sheets around 12 to 11 ka years ago. Interestingly, the Syltholm genome is entirely composed of WHG ancestry, suggesting that EHG ancestry did not reach southern Denmark in prehistory.

Furthermore, it is striking that the Syltholm genome also lacks Neolithic farmer ancestry, despite its relatively late date. This is all the more interesting because there is evidence for domesticated animals and farming practices at the site from around 6,000 BP (10). It suggests that admixture with farming populations might not have been as pervasive at the time. Having said that, the mtDNA genome we recovered from the resin belongs to haplogroup K1e, which is surprising, as it is more commonly associated with early farming communities (34–38). However, haplogroup K1 has also been observed in Mesolithic individuals in other parts of Europe (39) and it is possible, therefore, that it reached northern Europe during the Mesolithic. In any event, the lack of genome-wide Neolithic farmer ancestry in the Syltholm genome is worthy of note because it suggests that the genetic impact of early farming communities might not have been as extensive as once thought (34).

## Phenotypic traits

The phenotypic combination of blue eyes and otherwise dark hair and skin pigmentation has been previously noted in European hunter-gatherers, including La Braña (14) and Cheddar Man (40). Our results indicate that this phenotype was widespread in Mesolithic Europe and that the adaptive spread of light skin pigmentation in European populations only occurred later on in prehistory (32). The absence of adaptive alleles on the *MCM6* gene, which have consequences on lactase persistence into adulthood, has also been observed before (14) and fits well with the idea that the rise of dairy farming during the Neolithic created selective pressures that favoured the spread of alleles associated with lactose tolerance in adults (41). The fact that the Syltholm female carries derived alleles in a large number of genes that have a wide range of functions in the human immune system confirms earlier suggestions that the Neolithic transition, with its associated dietary changes and shifts in pathogen exposure, was not a major driving force of adaptive innovation relating to the immune response in modern humans (14).

## Oral microbiome

Previous studies (42–44) of microorganisms preserved in mineralised dental plaque (calculus) have shown changes to the ancestral oral microbiome over time. One of the most significant changes occurred with the Neolithic revolution. However, unlike dental calculus, which represents a long-term reservoir of the oral microbiota, the bacteria found in ancient mastics are more likely to give a snapshot of the species active at the time, and to reflect the microbiome of the oral mucosa instead of the plaque microbiome. As most modern clinical studies use saliva as sample material, this facilitates comparative studies. As seen in Fig. 3, the Syltholm sample has a similar microbial composition as modern oral microbiomes, but there are also some clear differences. Most notably, we observe an increase *Veillonellales* and a decrease in *Neisseriales* sp., which might indicate a dietary shift away from wild foods to a diet rich in carbohydrates and dairy products following the Neolithic transition (42).

## Diet

The Mesolithic/Neolithic transition in Denmark is generally thought to have been characterised by rapid and dramatic changes in subsistence strategies, from a marine diet in the Mesolithic, to a largely terrestrial-based diet in the Neolithic (23, 45). However, several lines of evidence suggest that freshwater and marine resources continued to be exploited during the Neolithic, including fishing equipment, faunal remains, and residue analyses (46–49). We identified several thousand reads in the resin that might be associated with diet, including reads mapping to mallard (*Anas platyrhynchos*) and eel (*Anguilla anguilla*) (Table 1). The presence of these two taxa in the sample suggests that the community at Syltholm continued to exploit wild resources well into the Neolithic. This is also supported by the presence of faunal remains at the site, such as duck (*Anas* sp.; NISP = 50), as well as hundreds of wooden leister prongs and bone points used for catching eel.

In summary, we demonstrate that chewed birch tar is an exceptional source of authentic ancient human and microbial DNA, presumably because of the hydrophobic and antibacterial properties of the resin. We recovered a complete ancient human genome (2.3×) and oral microbiota from one such artefact. While the human genome provides new information on the population dynamics of northern Europe at the time of the Neolithic transition, the microbial data shed light on the state of our ancestral oral microbiome. In addition, the human genome provides information on the evolution of skin pigmentation and other adaptive traits in humans. Lastly, the recovery of sequences derived from foodstuffs also provided insights into diet. This new source of ancient DNA opens up fascinating new possibilities for the field of ancient DNA, especially for time periods where human remains are scarce.

## Material and Methods

### Sample

The “gum” was found at site MLF906-II (Fig. S1) during excavations carried out by the Museum Lolland-Falster near Rødbyhavn, Denmark. The site spans the Mesolithic/Neolithic transition in Denmark (Fig. S2) and was heavily exploited during the Late Ertebølle and Early/Middle Neolithic periods. The piece of gum was found at a depth of c.2 m below surface in the north-eastern section of the site and measures c.3 cm in length and 1 cm in width (Fig. S3).

### Radiocarbon dating

The sample was only sparsely soluble in organic solvent, so we used an acid-base-acid (ABA) treatment. First, the sample was treated with 4% HCl (80°C) and then rinsed to neutrality with ultra-pure water. Second, a basic solution 1% NaOH (RT) was applied, and the reaction vessel rinsed again to neutrality. Finally, a further acid step was applied using 4% HCl (80°C) to ensure no atmospheric CO_2_ absorbed during the alkaline phase remained in the reaction vessel. After a last rinse to neutrality, the product was thoroughly air dried. An aliquot of approximately 4 mg was then weighed into a tin capsule for combustion in an Elemental Analyser (EA, IsotopeCube NCS, Elementar^®^). The EA was coupled to an Isotope Ratio Mass Spectrometer (IRMS, Isoprime^®^ 100), which allowed the δ^13^C value of the sample to be measured, as well as a fully automated cryogenic system that trapped the liberated CO_2_ into an airtight vessel. The vessel was manually transferred to a vacuum manifold, where a stoichiometric excess of H_2_(g) (1: 2.5) was added, and the sample CO_2_(g) reduced to graphite over an Fe(s) catalyst. The graphite was pressed into a cathode for radioisotope analysis in an accelerator mass spectrometer (MICADAS IonPlus^®^). The MICADAS generated an estimate of the ^14^C:^12^C ratio that was close to ±1‰, and from this data, and in accordance with all standard operations and conventions, the ^14^C date (in yrs BP) was calculated.

### FTIR and GC-MS analysis

The birch resin was analysed by Fourier Transform infrared (FTIR) spectroscopy and Gas Chromatography Mass Spectrometry (GC-MS). FTIR was carried out on a Perkin Elmer Spectrum 1000 FT-IR spectrometer using potassium bromide pellets (Fischer Scientific, IR Grade). GC-MS analysis was carried out on a Bruker SCION 456 GC-TQMS equipped with a Restek Rtx-5 capillary column (30 m, 0.25 mm ID, 0.25 μm) programmed for a 1 cm^3^ min^−1^ helium flow. 1 μl of sample was injected using a Programmed Temperature Vaporizer, held at 64°C for 0.5 min, raised to 315°C at 200°C min^−1^ and held at that temperature for 40 min. The split ratio was high from 0 to 0.5 min and then switched to 5. The GC oven temperature was held at 64°C for 0.5 min, then raised to 190°C at 10°C min^−1^ and then on to 315°C at 4°C min^−1^ and held at that temperature for 15 min. The sample was hydrolysed in methanolic KOH (Merck) and extracted with GC-grade tert-Butyl methyl ether (MTBE) after acidification. The MTBE extract was methylated with diazomethane (Sigma-Aldrich) prior to injection in the GC-MS system (50). The FTIR analysis produced a spectrum reminiscent of freshly produced birch resin (Fig. S4), while the GC-MS analysis revealed the presence of the triterpenes betulin and lupeol (Fig. S5), which are considered biomarkers of birch resin (51–55).

### Sample preparation and DNA extraction

We sampled c.250 mg from the gum and vortexed it briefly in 5% bleach solution to remove any surface contaminants. The sample was then washed in molecular grade water, left to dry, and cut into six smaller pieces weighing around 30-50 mg each. We then extracted DNA from the samples using three different methods: Samples 1-4 were incubated in 1 ml of a proteinase K-based digestion buffer on a rotor at 56°C for c.12 hrs. Subsequently, the supernatant for samples 1 and 2 was concentrated down to ~150 μl using Amicon Ultra centrifugal filters (MWCO 30 kDa), mixed 1:10 with a PB-based binding buffer (56), and purified using MinElute columns, eluting in 30 μl EB. For samples 3 and 4 we added an additional phenol-chloroform step, by adding 1 ml phenol (pH 8.0) to the digestion mix, followed by 1 ml chloroform:isoamyl alcohol. The supernatant was then concentrated and purified, as described above. Samples 5-6 were dissolved in 1 ml chloroform:isoamylalcohol without adding proteinase K. The chloroform:isoamylalcohol was mixed with 1 ml molecular grade water, that then was purified as described above. Following metagenomic profiling, the extracts prepared with chloroform were found to be contaminated with *Delftia* and *Methylophilus* sp., highlighting the danger of reagent and laboratory contamination for microbiome studies (57).

### Library preparation and sequencing

16 μl of each DNA extract were built into double-stranded libraries using the BEST protocol (58), with minor modifications as described in Mak et al. (59). One extraction NTC was included as well as a single library NTC. 10 μl of each library were amplified and indexed in 50 μl reactions, using a dual indexing approach, as described by Kircher et al. (60). The optimal number of PCR cycles was determined by qPCR (MxPro 3000, Agilent Technologies). The amplified libraries were purified using SPRI-beads and quantified on a 2200 TapeStation (Agilent Technologies) using High Sensitivity tapes. The amplified and indexed libraries were then pooled in equimolar amounts and sequenced on 1/10 of a lane of an Illumina HiSeq 2500 run in SR mode. Following initial screening, additional reads were obtained by pooling libraries #2, #3 and #4 in molar fractions of 0.2, 0.4 and 0.4, respectively and sequencing them on one full lane of an Illumina HiSeq 2500 run in SR mode.

### Decay rate estimate

To investigate the rate of human DNA degradation in the ancient chewing gum we examined the DNA read length distributions of the mapped reads, using a previously published method (61). The distribution follows a typical pattern of degraded DNA with an initial increase in the number of reads towards longer DNA fragments, followed by a decline. We observe that the declining part of the distribution follows an exponential decay curve (R^2^=0.99), as expected if the DNA had been randomly fragmented over time. Deagle et al. (62) showed that the decay constant (λ) in the exponential equation represents the fraction of broken bonds in the DNA strand (the damage fraction) and that 1/λ is the average theoretical fragment length in the DNA library. By solving the equation, we obtain a DNA damage fraction (λ) of 3.4%, which corresponds to a theoretical average fragment length (1/λ) of 29 bp (Table S3). We note that this is not directly comparable to the observed average length, which is affected by lab methods and sequencing technology. If the DNA is found in a stable matrix long term DNA fragmentation can be expressed as a rate and the damage fraction (λ, per site) can be converted to a decay rate (*k*, per site per year), when the age of the sample is known. By applying an estimated age of 5,700 years for the Syltholm resin, the corresponding DNA decay rate (*k*) is 5,96^−06^ breaks per bond per year, which corresponds to a molecular half-life of 1,162 years for a 100 bp DNA fragment. This means that after 1,162 years (post cell death), each 100 bp DNA stretch will have experienced one break on average. This estimated rate of DNA decay for the gum sample seems within the expected age for DNA preserved in a stable matrix in temperate climate zone. For example, the rate is close to that observed in the La Braña sample (14), preserved under similar temperatures as our gum sample (Table S3). By contrast, the DNA decay in human remains from warmer climates is much faster (63). Although these calculations are only based on a single sample, the results suggest that ancient mastics provide remarkable conditions for molecular preservation, with a rate of DNA decay that is comparable to that in well-preserved teeth or petrous bones.

### Data processing

Basecalling was done with bcl2fastq 2.2. Only sequences with correct indexes were kept. FastQ files were processed using PALEOMIX 1.2.12 (64). Adapters and low quality reads (Q<20) were removed using AdapterRemoval 2.2.0 (65) only retaining reads >25 bp. Trimmed and filtered reads were then mapped to hg19 (build 37.1) using BWA (66) with seed disabled to allow for better sensitivity (67), as well as filtering out unmapped reads. Only reads with a mapping quality ≥30 were kept and PCR duplicates were removed. MapDamage v.2.0.6 (68) was used to evaluate the authenticity of the retained reads as part of the PALEOMIX pipeline (64), using a subsample of 100k reads per sample (Fig. S5). For the population genomic analyses, we merged the ancient sample with individuals from the Human Origin dataset (13) and over >200 other previously published ancient genomes. At each SNP in the human origin dataset, we sampled the allele with more reads in the ancient sample, resolving ties randomly, resulting in a pseudohaploid ancient sample.

### MtDNA and contamination estimates

We used Schmutzi (69) to determine the endogenous consensus mtDNA sequences and to estimate the amount of modern human contamination. Reads were mapped to the Cambridge reference sequence (rCRS) and filtered for MAPQ ≥30. Haploid variants were called using the endoCaller program implemented in Schmutzi (69) and only variants with a posterior probability exceeding 50 on the PHRED scale (probability of error: 1/100000) were retained. We then used Haplogrep 2.2 (70) to determine the mtDNA haplogroups, specifying PhyloTree (build 17) as the reference phylogeny (71).

### Imputation and phenotyping

We used information from 41 SNPs from the HirisPlex-S system (17) to identify phenotypes such as skin, eye and hair colour in the ancient individual. First, we used angsd (72) to obtain genotype likelihoods in a 5 Mb region around these SNPs. Subsequently, we used the 5 Mb region to estimate the underlying haplotypes carried by the Syltholm individual at these loci, and from these haplotypes imputed the genotypes at the 41 SNPs of interest. We used multiple posterior probability thresholds, ranging from 0.95 to 0.50, to filter the imputed genotypes. These imputed genotypes were uploaded to the HIrisPlex-S website to obtain the predicted outcomes for the pigmentation phenotypes. In addition, using the same steps as outlined above, we imputed 82 additional SNPs, known to play a role in lactase persistence into adulthood, and immune response (14). We used the imputed genotypes to check for the presence of derived alleles.

### Principal component analysis

Principal component analysis was performed using smartPCA (73) by projecting the ancient individuals onto a modern reference panel including around 1,000 modern Eurasian individuals from the HO dataset (13) using the option lsq project. Prior to performing the PCA the data set was filtered for a minimum allele frequency of at least 5% and a missingness per marker of at most 50%. To mitigate the effect of linkage disequilibrium, the data were pruned in a 50-SNP sliding window, advanced by 10 SNPs, and removing sites with an R^2^ larger than 0, resulting in a final data set of 593,102 SNPs.

### Model-based clustering

We used ADMIXTURE v.1.3.0 (15) to estimate the admixture proportions of the ancient individuals in our merged dataset. The algorithm was run in supervised mode, specifying typical Western hunter-gatherers (Loschbour), Eastern hunter-gatherers (Samara), and Neolithic farmers (Barcın), as potential ancestral source populations. To help visualise the results, we plotted population averages for a selected number of ancient populations (Fig. 2.b).

### D- and f-statistics

*D*- and *f*-statistics were computed using *AdmixTools* (74). To estimate the amount of shared drift between a set of ancient genomes, including the Syltholm genome, WHG, and EHG, we calculated *f*_4_-statistics of the form *f*_4_(Yoruba, *X*; EHG, WHG), where *X* stands for one of 15 ancient genomes, including the Syltholm genome. Standard errors were calculated using a weighted block jackknife. To confirm the absence of EHG gene flow in the Syltholm genome and to contrast this result with those obtained for other Scandinavian hunter-gatherers, we calculated a set of *D*-statistics of the form *D*(Yoruba, EHG; WHG, X) where *X* stands for one of 15 ancient genomes, including the Syltholm genome. Lastly, we used the *qpAdm* algorithm from the *AdmixTools* package (74) to model different admixture scenarios and estimate admixture proportions in the Syltholm genome based on *f*_4_-statistics and assuming WHG, EHG, and Neolithic farmers (Barcın) as potential ancestral source (“left”) populations and one modern (Dinka) and seven ancient genomes, including the Mal’ta genome (75), as outgroup (“right”) populations.

### Metagenomic profiling

We used MetaPhlan2 (76) to create a metagenomic profile based on the non-human reads. The reads were first aligned to the MetaPhlan2 database (76, 77) using Bowtie2 v2.2.9 aligner (78). PCR duplicates were removed using PALEOMIX filteruniquebam (67). For cross tissue comparisons 689 human microbiome profiles published in the Human Microbiome Project Consortium (79) were initially used, comprising samples from the mouth (N=382), skin (N=26), gastrointestinal tract (N=138), urogenital tract (N=56), airways and nose (N=87). Pairwise ecological distances amongst the profiles were computed at genera and species level using taxon relative abundances and the vegdist function from the vegan package (http://cran.r-project.org/package=vegan). These were used for principal coordinate analysis (PCoA) of Bray-Curtis distances with the R function pcoa. Subsequently, we calculated the average relative abundance of each genus for each of the body sites present in the Human Microbiome Project and visualised the abundance of microbial orders of our sample and the HMP body sites with MEGAN6 (80). To further characterise the metagenomic reads we used MGmapper v.2.7 (18). Trimmed and filtered sequences were split in ten FastQ files, and MGmapper was used on best mode to first align these to the PhiX database, and then align the non-PhiX reads against the human, viral, plant, mammal, other vertebrates, invertebrates, human oral microbiome, and bacterial database, where only unmapped reads were aligned to the next database. The results of the ten read subsets were then concatenated and processed with MGmapper_classify. Besides MGmapper’s default filters, hits with a lower ratio of unique reads/all reads than 0.01 were excluded. To validate the assignments we then used a reference-based mapping approach. Human reads were removed by first aligning all trimmed and filtered reads to the human reference genome hg19 (build 37.1) using BWA and subsequently extracting all unmapped reads. The reference genomes of taxa of interest were obtained from the NCBI Reference Sequence Database. The trimmed and filtered non-human sequences were aligned to each reference genome with BWA (66) and PCR duplicates removed using Picard Tools v.2.13.2 (http://broadinstitute.github.io/picard/). Sequences with mapping quality below 35 were removed, and mapDamage v.2.0.6 (81) was used to estimate postmortem damage (Fig. S7). The breadth and depth of coverage (Fig. S8) were then calculated with bedtools v.2.27.1 (82) and visualized with circos v.0.69-6 (83). As a final check, sequences that aligned to the reference genome were extracted with bamtools v.2.5.1 (84) and subsequently assembled with MEGAHIT v.1.1.1 (85). The resulting assemblies were then used as a query for a nucleotide BLAST search (86) against the nr/nt database.

## Acknowledgments

We thank the Museum Lolland-Falster for access to the sample and Miren Iraeta Orbegozo for technical assistance.

## Author contributions

TZTJ and HS designed and led the study. SAS provided the sample for analysis. TZTJ, MHSS, MRE, MCC, MNM and MWD generated the data. TZTJ, JN, KHI, AKF, SG, SHN, MEA, LTL, AJT, MCC, MNM and HS analyzed the data. TZTJ, JN, LTL, AJT, SAS, MJC, MTPG, MS, SR and HS interpreted the data. TZTJ, JN and HS wrote the manuscript with input from all the other authors.

## Data and materials availability

Alignment data are available through the European Nucleotide Archive under accession number PRJEB30280.

---

**Fig. S1.**
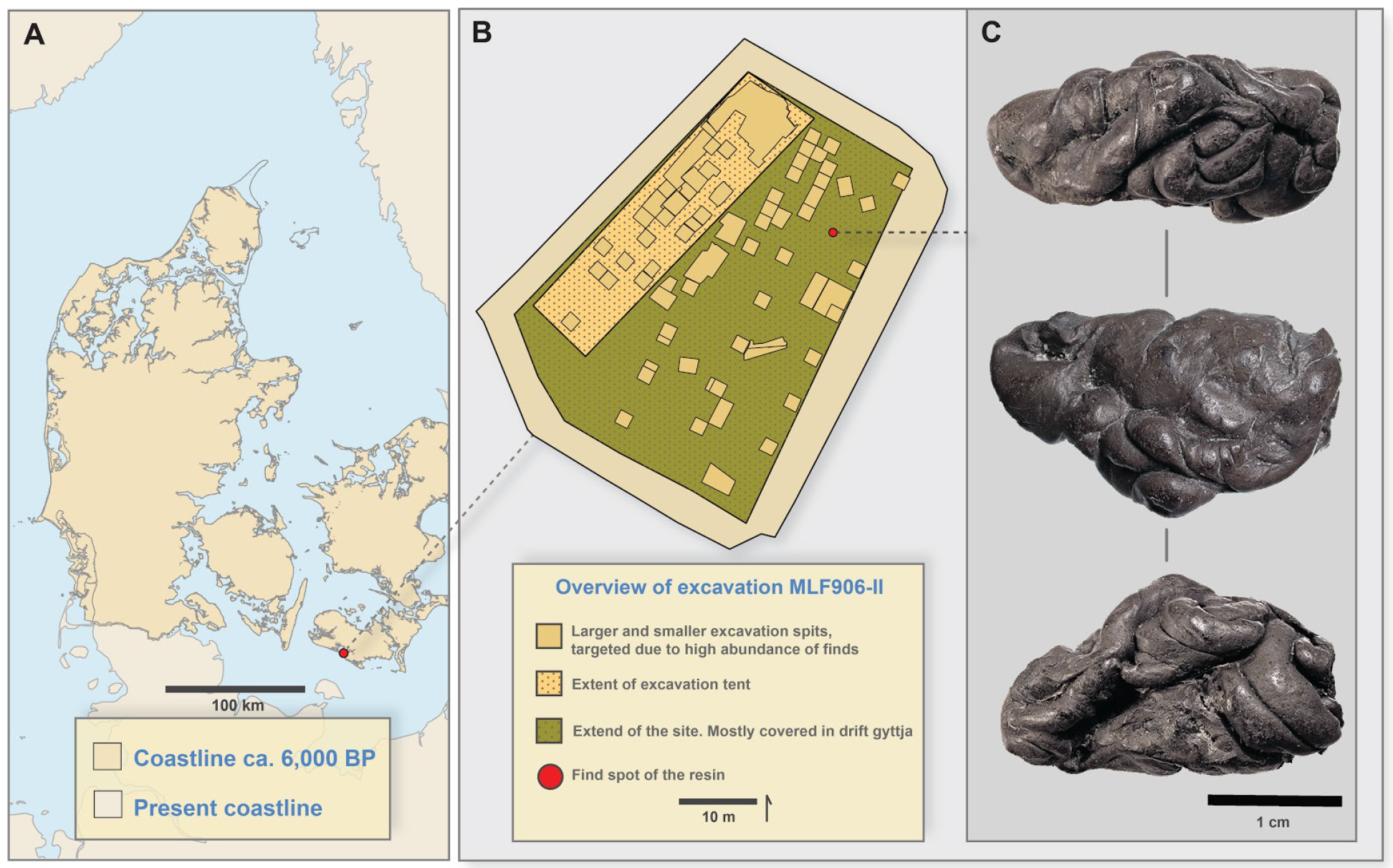
**a)** Overview of Denmark with approximate coastline during the Littorina sea and the findspot at Syltholm, Rødbyhavn, Denmark (Danish coastline 6.000 bp after (87)). **b)** GIS digitisation of the excavation and findspot of the resin, **c)** photo of the birch resin.

**Fig. S2.**
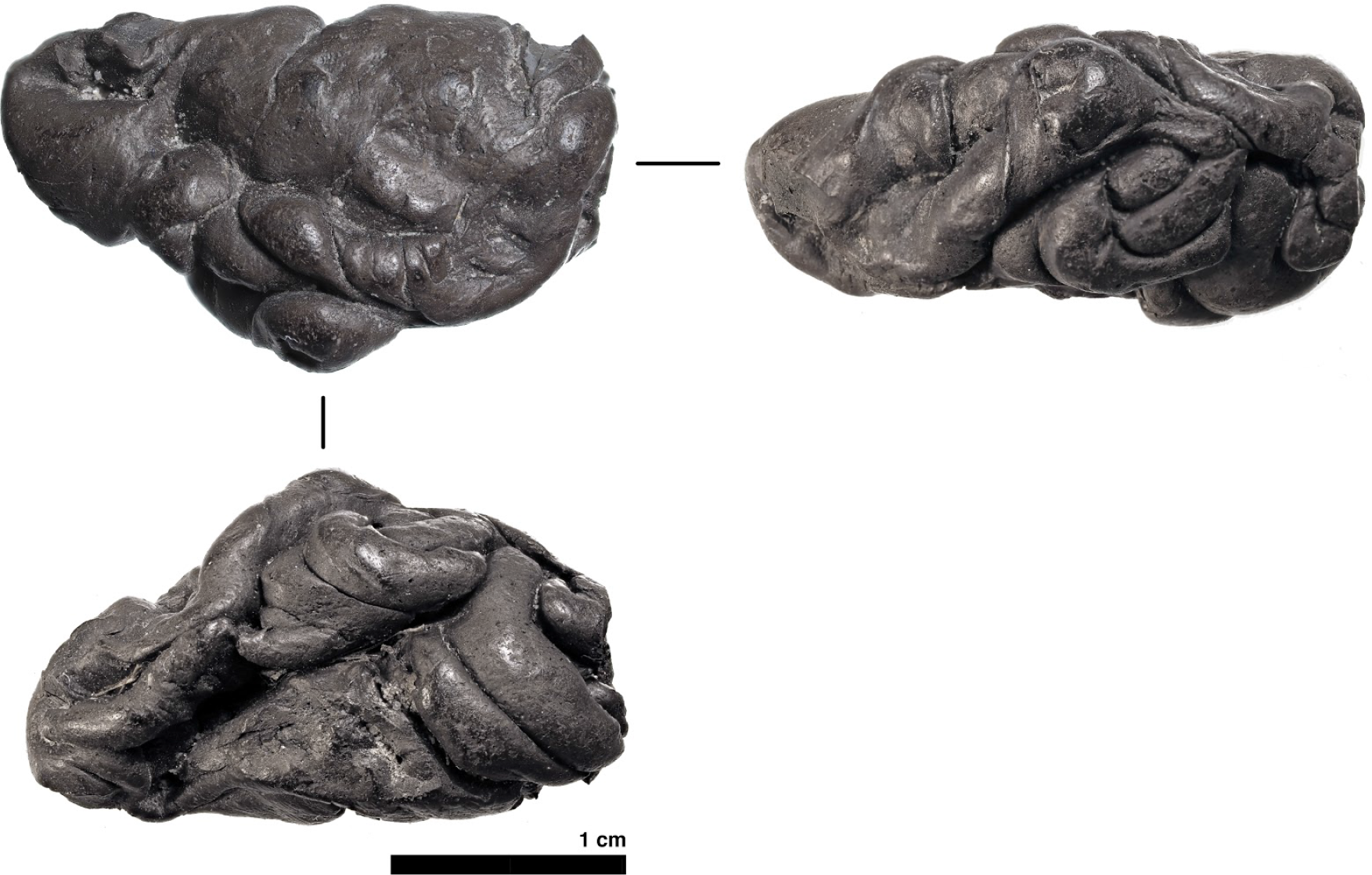
Close-up photograph of the Syltholm resin.

**Fig. S3.**
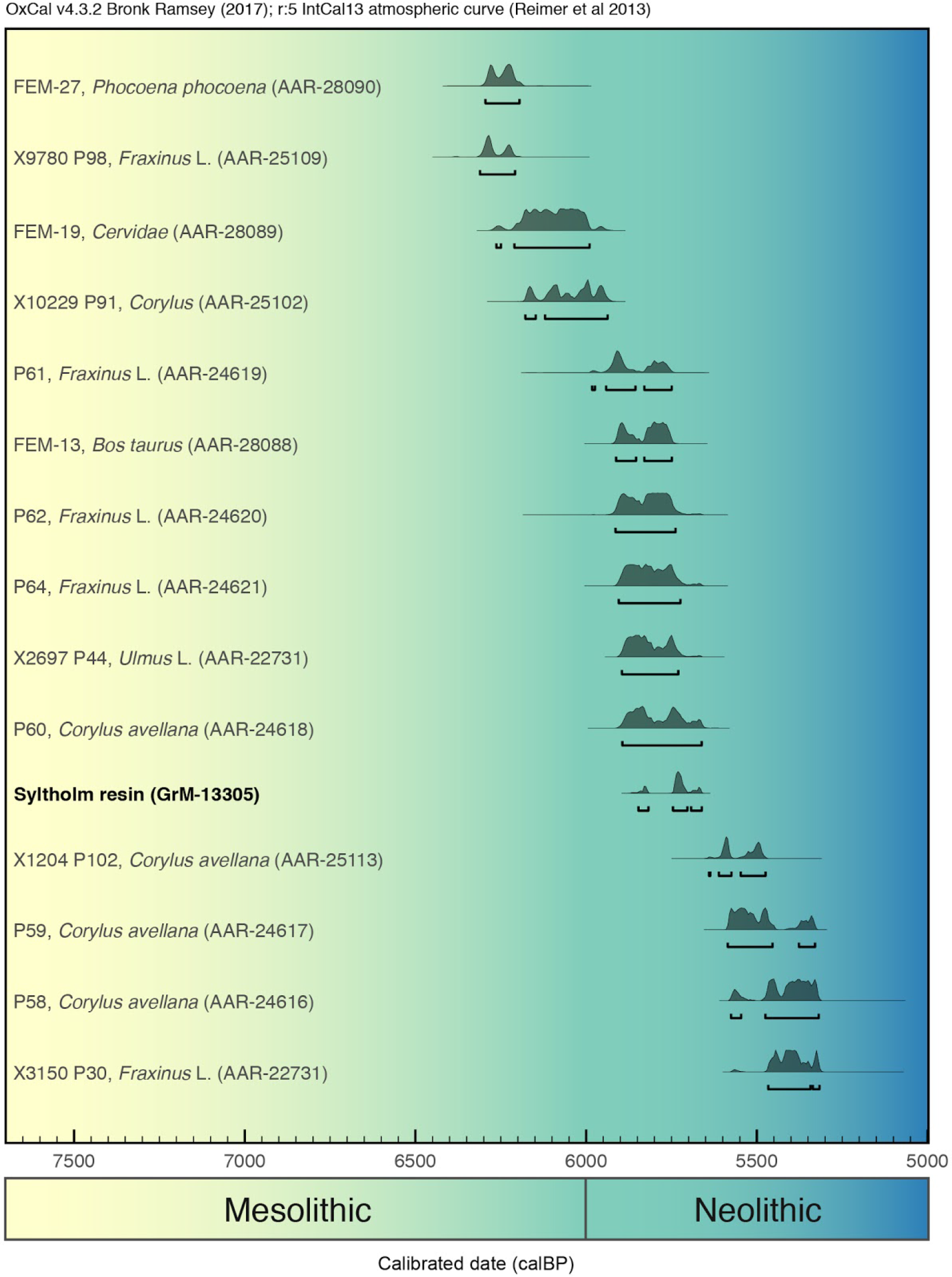
Radiocarbon chronology for Syltholm site MLF906-II based on a series of 15 calibrated radiocarbon dates, including the “gum”.

**Fig. S4.**
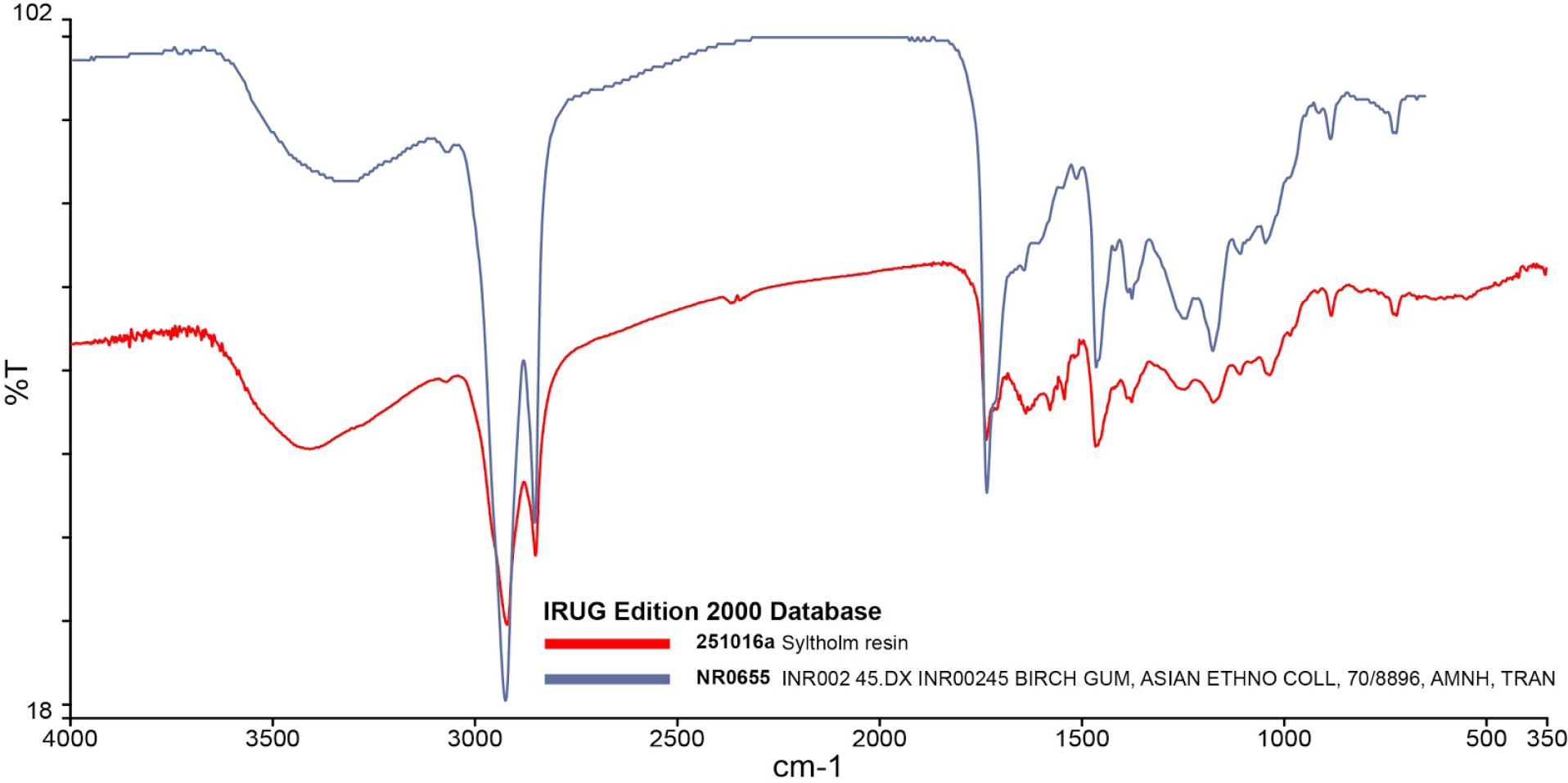
FT-IR spectra of the Syltholm resin and a modern birch sample.

**Fig. S5.**
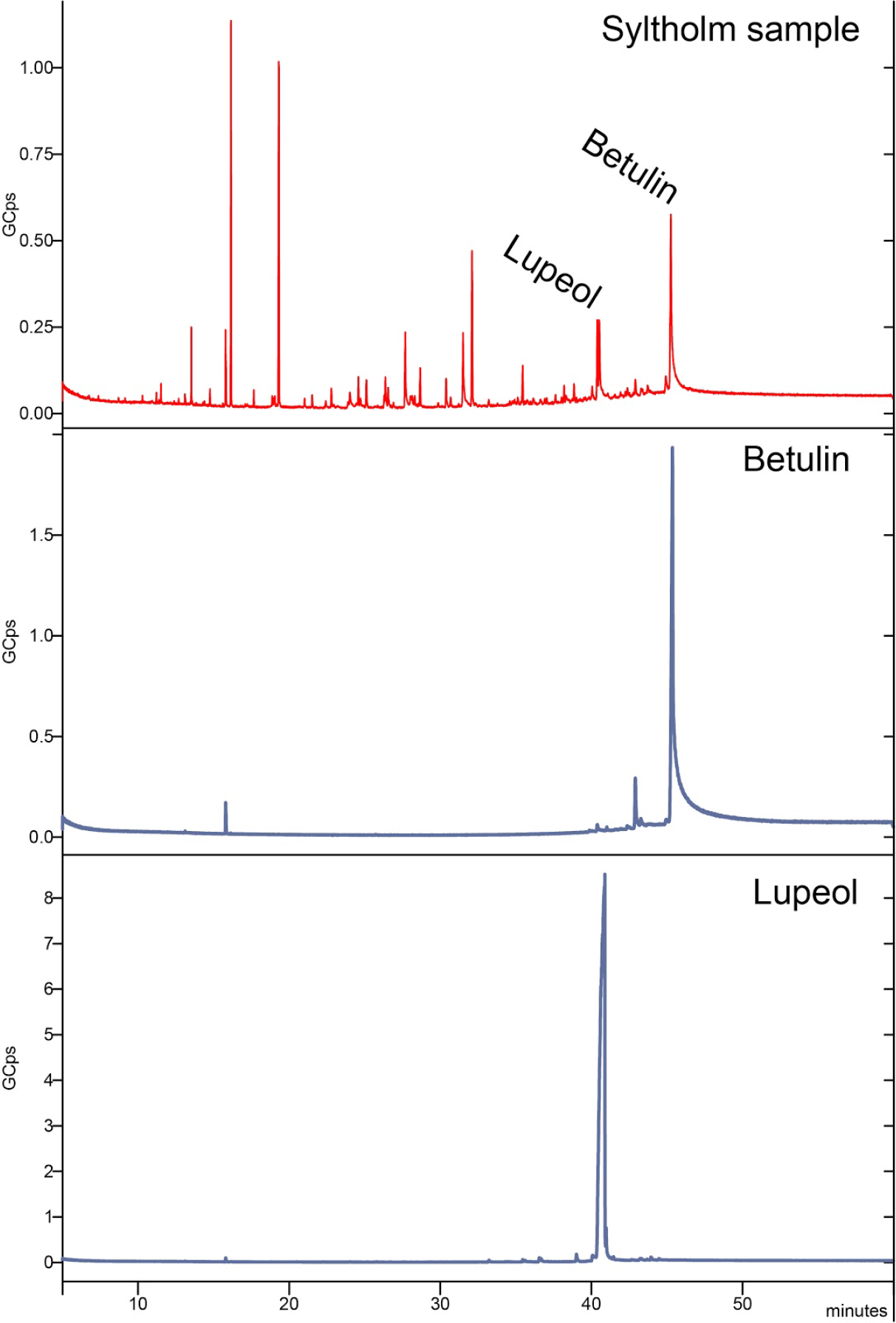
GC-MS plot of the Syltholm sample (top), pure betulin reference (middle) and pure lupeol reference (bottom).

**Fig. S6.**
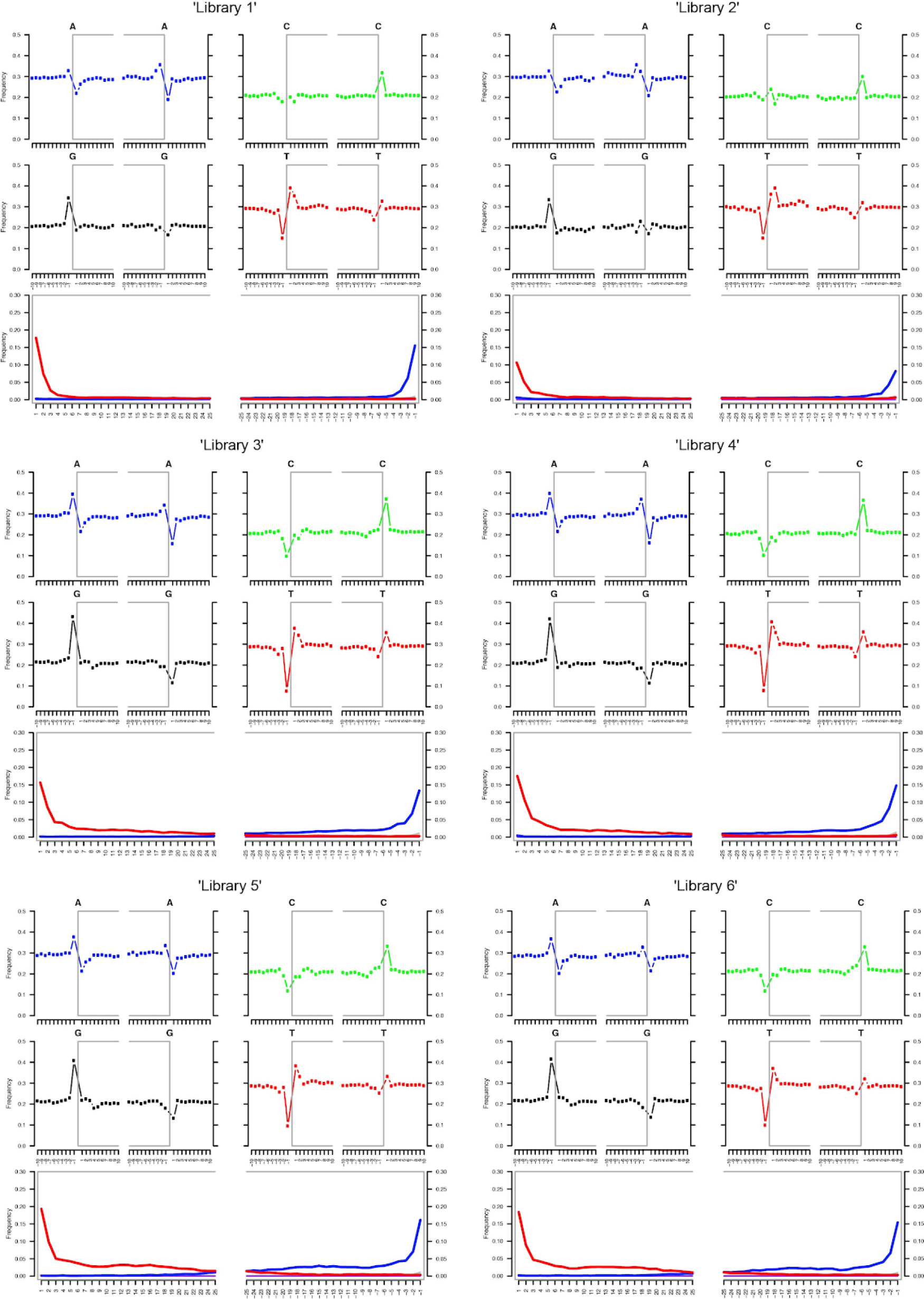
MapDamage (88) plots for reads mapping to the human reference genome (hg19), by library.

**Fig. S7.**
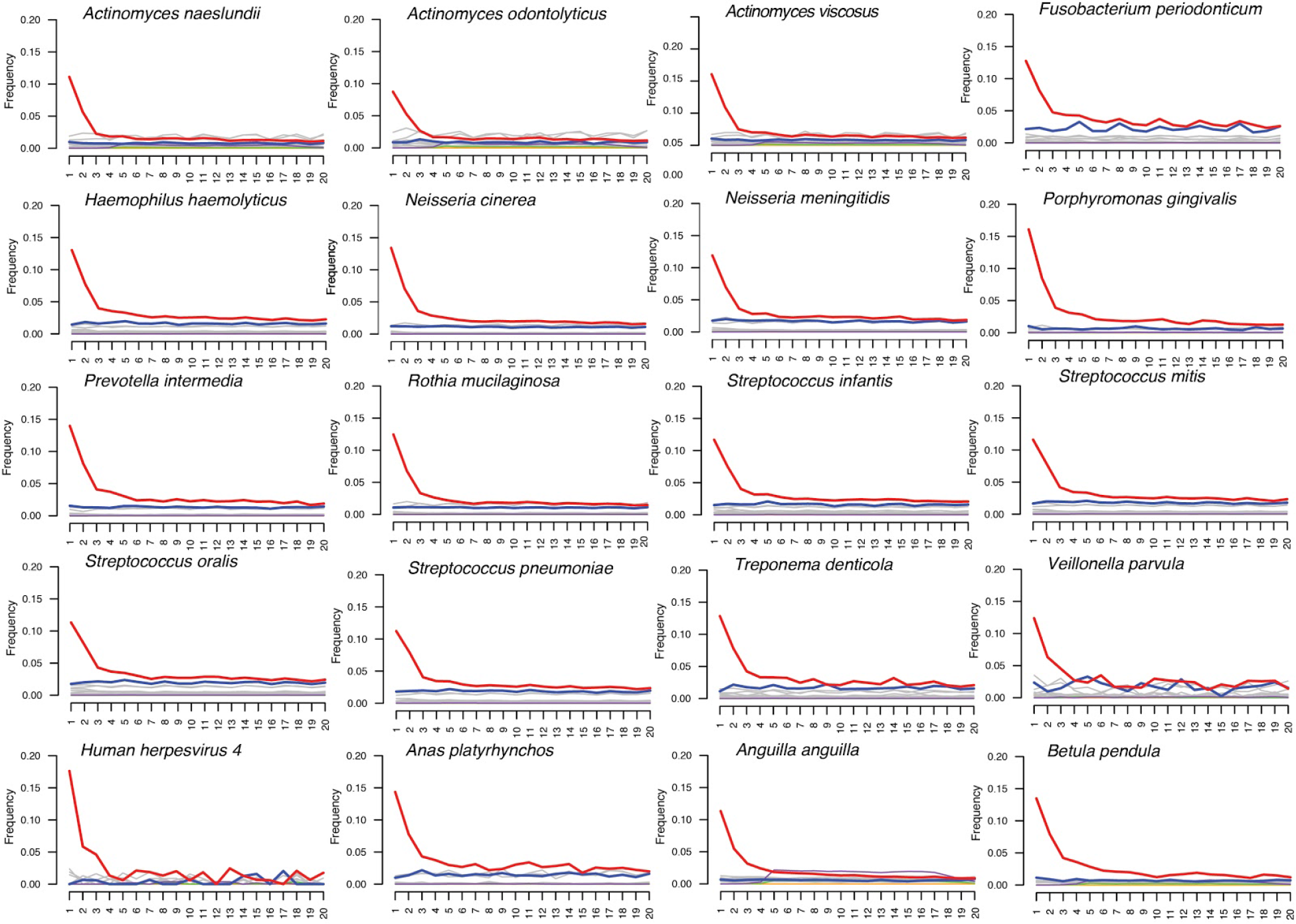
MapDamage (88) plots for non-human taxa recovered from the Syltholm resin.

**Fig. S8.**
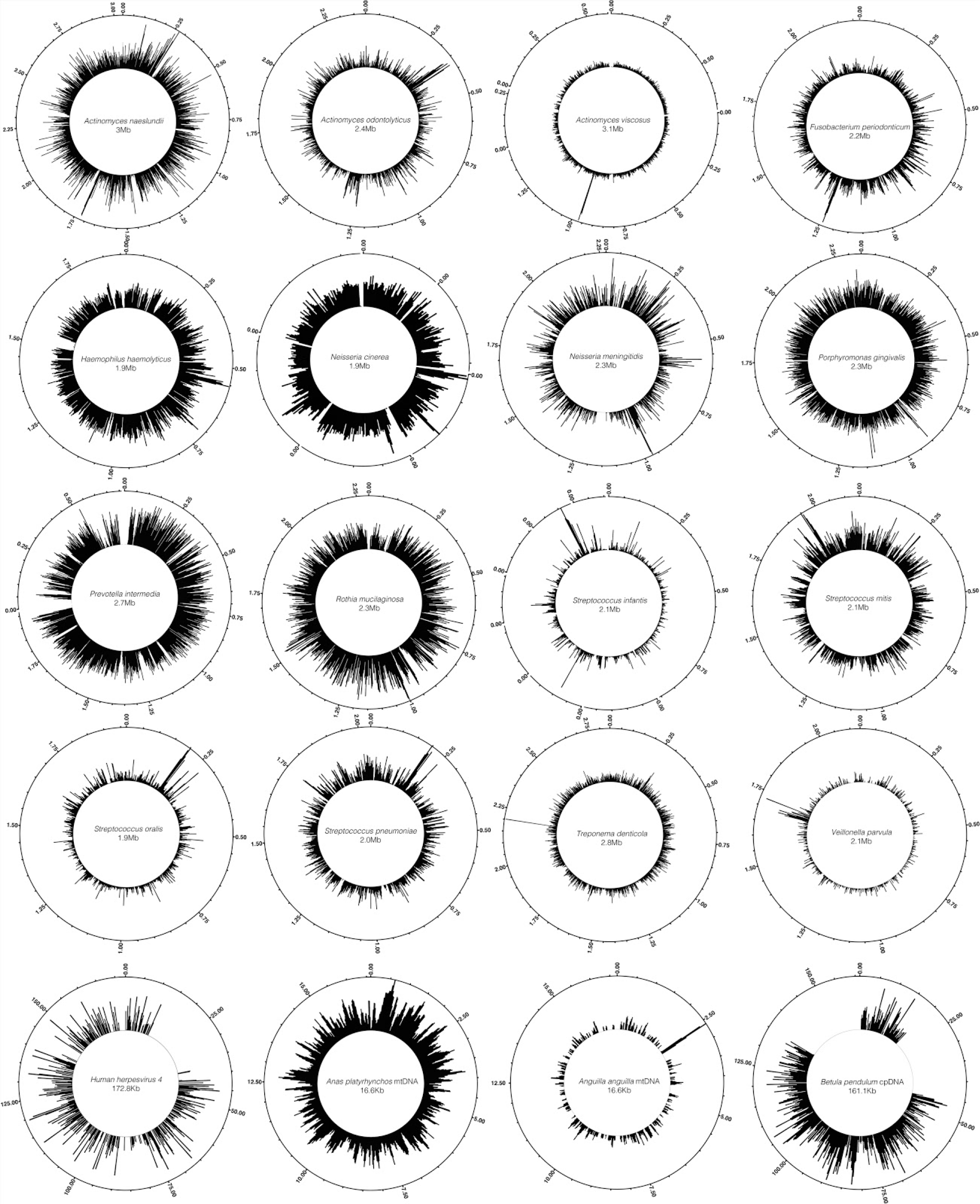
Coverage plots for non-human taxa recovered from the Syltholm resin.

**Table S1.**
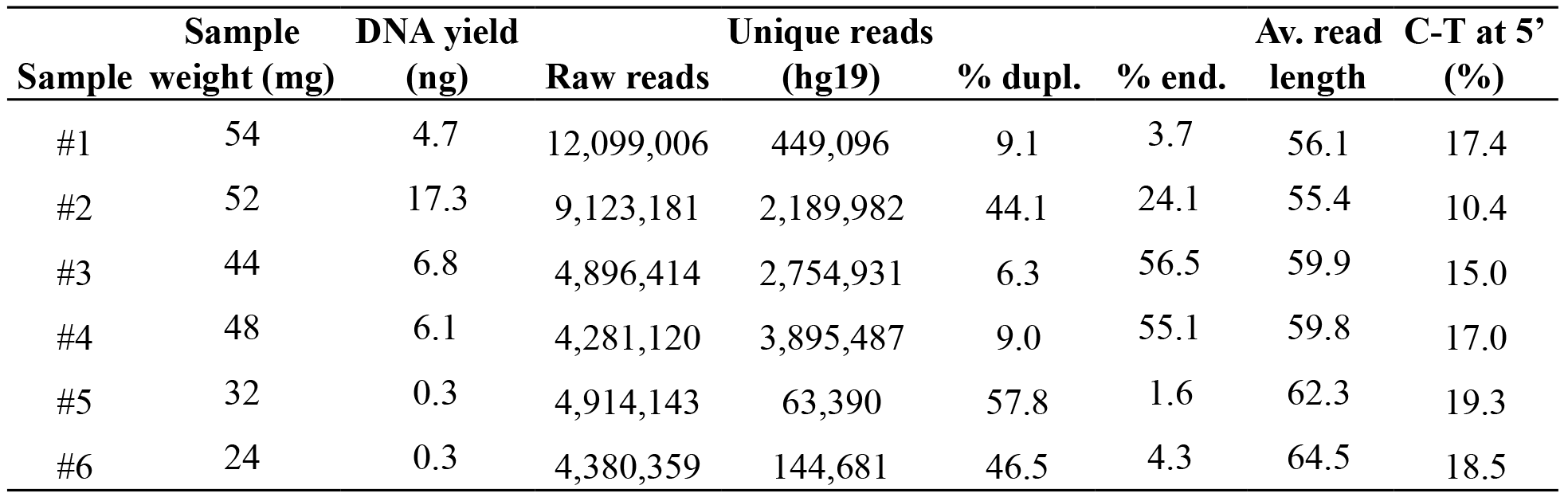
Screening results for the Syltholm resin. Total DNA yields (ng) were measured on raw extracts using the TapeStation. Analysis ready reads refer to reads that could be mapped to the human reference genome (hg19) after removing duplicates. Deamination rates at 5’ ends of DNA fragments were determined using MapDamage 2.0 (88).

**Table S2.** Deep-sequencing results for the Syltholm resin. Analysis ready reads refer to reads that could be mapped to the human reference genome (hg19) after removing duplicates. Deamination rates at 5’ ends of DNA fragments were determined using MapDamage 2.0 (88).

| **Total reads** | **Unique reads**<br>**(hg19)** | **% dupl.** | **% end.** | **Av. read**<br>**length** | **C-T at 5'**<br>**(%)** | **mtDNA**<br>**cont. (%)** | **>1X (%)** | **DoC** |
| --- | --- | --- | --- | --- | --- | --- | --- | --- |
| 389,930,326 | 120,585,267 | 42.7 | 31.2 | 59.9 | 16.2 | 1-3 | 78.9 | 2.3 |

**Table S3.**
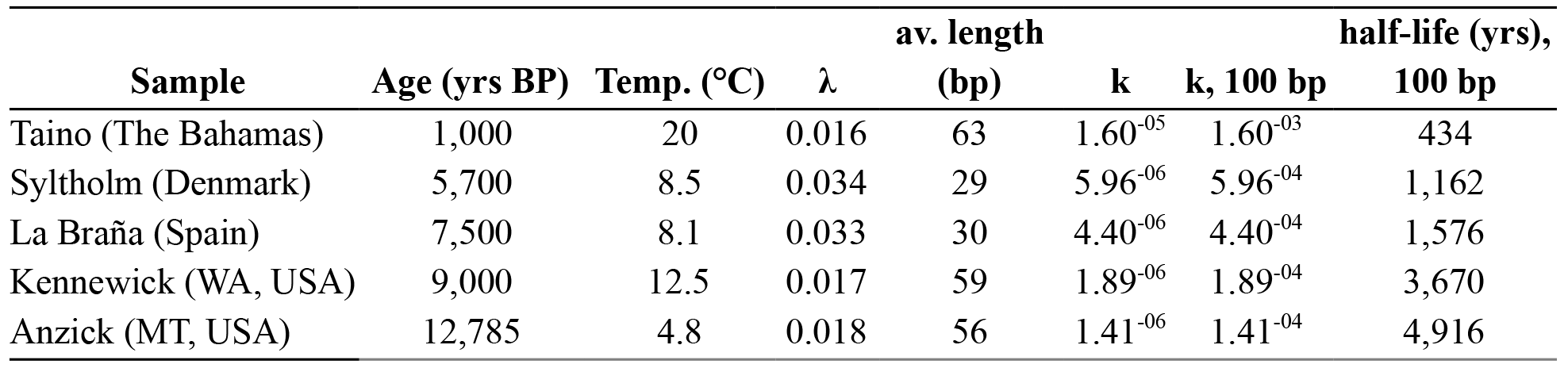
Molecular decay rates for the Sylthom genome and other previously published ancient genomes from different contexts (14, 63, 89, 90).

**Table S4.** *f*_4_-statistics of the form *f*_4_(Yoruba, *X*; EHG, WHG) estimating the amount of shared drift between the ancient genomes (*X*), EHG and WHG.

| Pop2 (X) | $f_4$-stat | SE | Z | BABA | ABBA | SNPs |
| --- | --- | --- | --- | --- | --- | --- |
| Syltholm | 0.011917 | 0.000698 | 17.063 | 6281 | 4903 | 115687 |
| La Braña | 0.012022 | 0.000525 | 22.894 | 29695 | 23219 | 538716 |
| Hum1 | -0.001431 | 0.000644 | -2.224 | 9966 | 10262 | 207167 |
| Hum2 | -0.001152 | 0.000592 | -1.947 | 26029 | 26646 | 536119 |
| Steigen | -0.001494 | 0.000565 | -2.645 | 20418 | 21047 | 421170 |
| Motala1 | 0.00216 | 0.000575 | 3.755 | 17719 | 16950 | 355954 |
| Motala2 | 0.003681 | 0.000546 | 6.747 | 22397 | 20771 | 441690 |
| Motala3 | 0.002856 | 0.000529 | 5.396 | 13191 | 12427 | 267396 |
| Motala4 | 0.003171 | 0.000578 | 5.484 | 22361 | 20955 | 443456 |
| Motala6 | 0.002229 | 0.000554 | 4.023 | 18922 | 18073 | 380891 |
| Motala12 | 0.002848 | 0.000545 | 5.223 | 25448 | 24005 | 506761 |

**Table S5.** *D*-statistics of the form *D*(Yoruba, *EHG*; WHG, *X*) testing for EHG gene flow in the Syltholm genome.

| **Pop4 (Z)** | **D-stat** | **SE** | **Z** | **BABA** | **ABBA** | **SNPs** |
| --- | --- | --- | --- | --- | --- | --- |
| Syltholm | 0.0179 | 0.006957 | 2.575 | 4965 | 4790 | 115800 |
| Hum1 | 0.046 | 0.006078 | 7.562 | 10345 | 9436 | 207431 |
| Hum2 | 0.055 | 0.00517 | 10.64 | 26829 | 24032 | 536745 |
| Steigen | 0.0612 | 0.005076 | 12.064 | 21173 | 18729 | 421659 |
| Motala1 | 0.0467 | 0.005117 | 9.134 | 17104 | 15576 | 356117 |
| Motala2 | 0.0479 | 0.004969 | 9.638 | 20965 | 19049 | 441955 |
| Motala3 | 0.0401 | 0.005147 | 7.8 | 12536 | 11569 | 267482 |
| Motala4 | 0.0443 | 0.005111 | 8.675 | 21113 | 19321 | 443722 |
| Motala6 | 0.0528 | 0.004972 | 10.625 | 18202 | 16375 | 381076 |
| Motala12 | 0.0478 | 0.004786 | 9.984 | 24237 | 22026 | 507157 |

**Table S6.** Chi-square statistics for different *qpAdm* models, assuming WHG, EHG, and Neolithic farmers (Barcın) as potential ancestral source (“left”) populations and one modern (Dinka) and seven ancient genomes, including the Mal’ta genome (75), as outgroup (“right”) populations.

| **fixed pat** | **wt** | **dof** | **chisq** |
| --- | --- | --- | --- |
| 000 (EHG, WHG, NEO) | 0 | 6 | 3.086 |
| 001 (EHG, WHG) | 1 | 7 | 5.936 |
| 010 (EHG, NEO) | 1 | 7 | 39.322 |
| 100 (WHG, NEO) | 1 | 7 | 6.152 |
| 011 (EHG) | 2 | 8 | 126.612 |
| 101 (WHG) | 2 | 8 | 9.15 |
| 110 (NEO) | 2 | 8 | 58.597 |

